# Rapid dissemination and monopolization of viral populations in mice revealed using a panel of barcoded viruses

**DOI:** 10.1101/769976

**Authors:** Broc T. McCune, Matthew R. Lanahan, Benjamin R. tenOever, Julie K. Pfeiffer

## Abstract

The gastrointestinal tract presents a formidable barrier for pathogens to initiate infection. Despite this barrier, enteroviruses, including coxsackievirus B3 (CVB3), successfully penetrate the intestine to initiate infection and spread systemically prior to shedding in stool. However, the effect of the gastrointestinal barrier on CVB3 population dynamics is relatively unexplored, nor are the selective pressures acting on CVB3 in the intestine well-characterized. To examine viral population dynamics in orally infected mice, we produced over one hundred CVB3 viruses harboring unique nine nucleotide “barcodes.” Using this collection of barcoded viruses, we found diverse viral populations throughout each mouse within the first day post-infection, but by 48 hours the viral populations were dominated by less than three barcoded viruses in intestinal and extra-intestinal tissues. Using light-sensitive viruses to track replication status, we found diverse viruses had replicated prior to loss of diversity. Sequencing whole viral genomes from samples later in infection did not reveal detectable viral adaptations. Surprisingly, orally inoculated CVB3 was detectable in pancreas and liver as soon as 20 minutes post inoculation, indicating rapid systemic dissemination. These results suggest rapid dissemination of diverse viral populations, followed by a major restriction in population diversity and monopolization in all examined tissues. These results underscore a complex dynamic between dissemination and clearance for an enteric virus.

**Importance:** Enteric viruses initiate infection in the gastrointestinal tract but can disseminate to systemic sites. However, the dynamics of viral dissemination are unclear. In this study, we created a library of 135 barcoded coxsackieviruses to examine viral population diversity across time and space following oral inoculation of mice. Overall, we found that the broad population of viruses disseminates early, followed by monopolization of mouse tissues with three or fewer pool members at later time points. Interestingly, we detected virus in systemic tissues such as pancreas and liver just 20 minutes post-oral inoculation. These results suggest rapid dissemination of diverse viral populations, followed by a major restriction in population diversity and monopolization in all examined tissues.

## Introduction

RNA virus populations are dynamic. Population diversity arises through mutations introduced by the viral RNA-dependent RNA polymerases, which lack proofreading. Although many mutations are detrimental, viruses can benefit from diversity within viral populations since diversity may facilitate replication and dissemination within hosts (1,2). Viral diversity can be reduced through selective pressures or stochastic events. Because viral diversity is a fundamental parameter in viral evolution and virulence, understanding population dynamics and the factors that influence viral populations is a crucial question, with consequences for emergence of new pathogens, evolution of new traits, and vaccine design.

Previous studies have examined viral population dynamics in a variety of systems. Changes in viral populations can be quantified using genetically marked viruses encoding short unique nucleic acid sequences, called “barcodes” (3–7). Several of these studies revealed founder effects, an extreme reduction in viral population diversity in which a small sub-population of viruses invade a new tissue or host. This has been described for viruses of plants (8) and animals, including HIV (9), HCV (10), Zika (11,12), influenza (3,13), and poliovirus (6). The gastrointestinal (GI) tract restricts dissemination of lumenal contents, including enteric viruses. Previous work in our lab demonstrated that the GI tract limits dissemination of the enteric virus poliovirus, thus reducing the diversity of viral populations in a variety of tissues after oral inoculation (6). However, these studies lacked detailed temporal analysis of viral populations, so how the GI tract influences enteric viral populations over time is unknown.

Coxsackievirus B3 (CVB3) spreads by the fecal-oral route. After oral infection and passage through the GI tract, CVB3 disseminates to and replicates in extra-intestinal tissues prior to shedding in stool. Immune competent mice are orally susceptible to CVB3 infection, but do not succumb to disease. Immune deficient mice lacking the *interferon alpha/beta receptor* (*Ifnar-/-*) support high levels of viral replication and succumb to disease, providing a more tractable model to study infection with enteroviruses by the natural oral route (14–17).

In this study, we examined the dissemination patterns of CVB3 over time to assess how viral diversity changes across different tissues. Using a collection of barcoded viruses, we found diverse viral populations across many tissues within the first day after oral inoculation of mice. Surprisingly, dissemination was extremely rapid, with diverse viral populations present in systemic sites at 20 minutes post-infection. However, by 48 hours post-infection, a small number of viruses monopolized the entire mouse, without detectable virus adaptation. These results reveal dynamic viral dissemination and clearance for an enteric virus.

## Results

### Multiple unique viruses from oral inoculum reach extra-intestinal tissues

We created genetically tagged viruses that allow discrimination of individual members of the population to monitor viral population dynamics. We engineered 135 CVB3 viruses encoding unique 9 nucleotide “barcodes” in the 5’ untranslated region (nucleotide 709, Fig. 1A, B). Insertion of a barcode sequence only marginally slowed viral replication in HeLa cells (Fig. 1C), suggesting that barcoded viruses have high enough fitness for *in vivo* experiments. Furthermore, after passaging for 10 viral replication cycles in HeLa cells, the viral barcode was still present, indicating the barcode insertion was well tolerated (data not shown). Each of the 135 barcoded viruses were generated individually prior to combining equal titers of each to create the barcode virus library. Sequencing replicates of the 135-member barcode virus library confirmed approximately equal representation of each pool member and that the sequencing procedure was reproducible (Fig. 1D).

**Figure 1.**
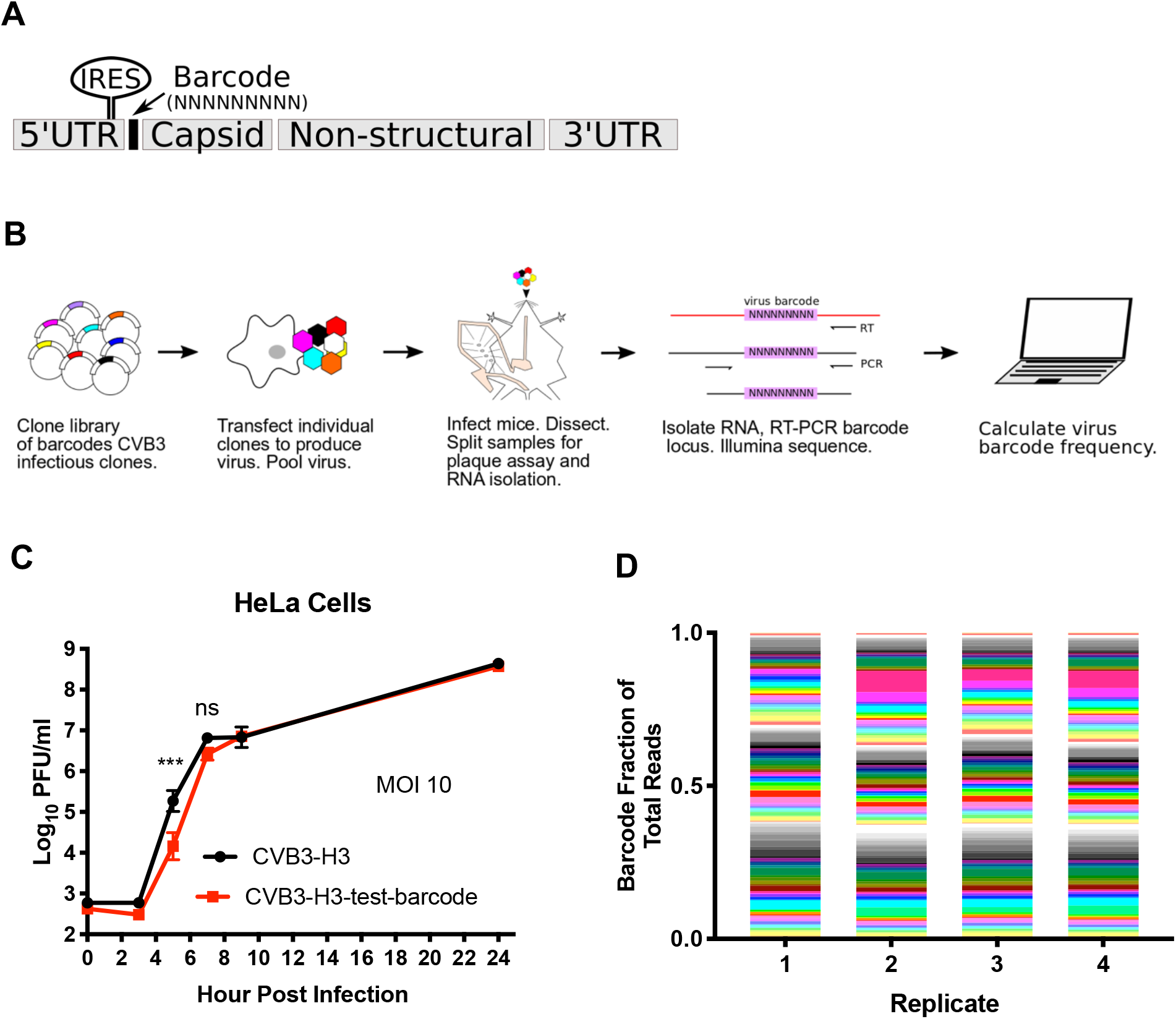
Construction of barcoded collection of CVB3 viruses. A. Random 9 nucleotide sequence was inserted into CVB3 5’UTR (nucleotide 709) between the internal ribosome entry site (IRES) and start of coding sequence. B. Workflow for production and analysis of virus-barcode frequencies in 135 CVB3 barcode virus pool. Different colors in plasmids and virions represent uniquely barcoded viruses. C. Growth curve of a representative single barcode CVB3 (barcode ATCGTACCA) and WT in HeLa cells (n=4, 2-way ANOVA, Sidak post-test, SEM shown). D. Sequencing ratios for each barcode in 4 cDNA replicates of CVB3-barcode virus stock.

To quantify how viral populations change over time and space, we infected *Ifnar-/-* mice orally with 1×10^9^ PFU of the 135-member CVB3 library and collected stool and tissues at various time points. At time points within the first day of infection, 7.5 and 19 hours post infection (hpi), we detected all 135 viruses in the upper GI tract (Fig. 2), indicating relatively rapid dissemination of the broad viral population in the inoculum. Furthermore, these results demonstrate that our sequencing and analysis pipeline was sensitive and unbiased for use with mouse tissues. Interestingly, at 48 and 72 hpi, a small number of pool members, generally three or less, comprised the majority of viral barcodes in each tissue (Fig. 2). Furthermore, nearly all tissues within an animal contained the same small group of viral barcodes, suggesting a relatively unencumbered flow of virus between tissues after 19hpi. These high-frequency viruses were unique to each mouse, indicating that no barcoded virus was uniquely fit to replicate and disseminate in mice.

**Figure 2.**
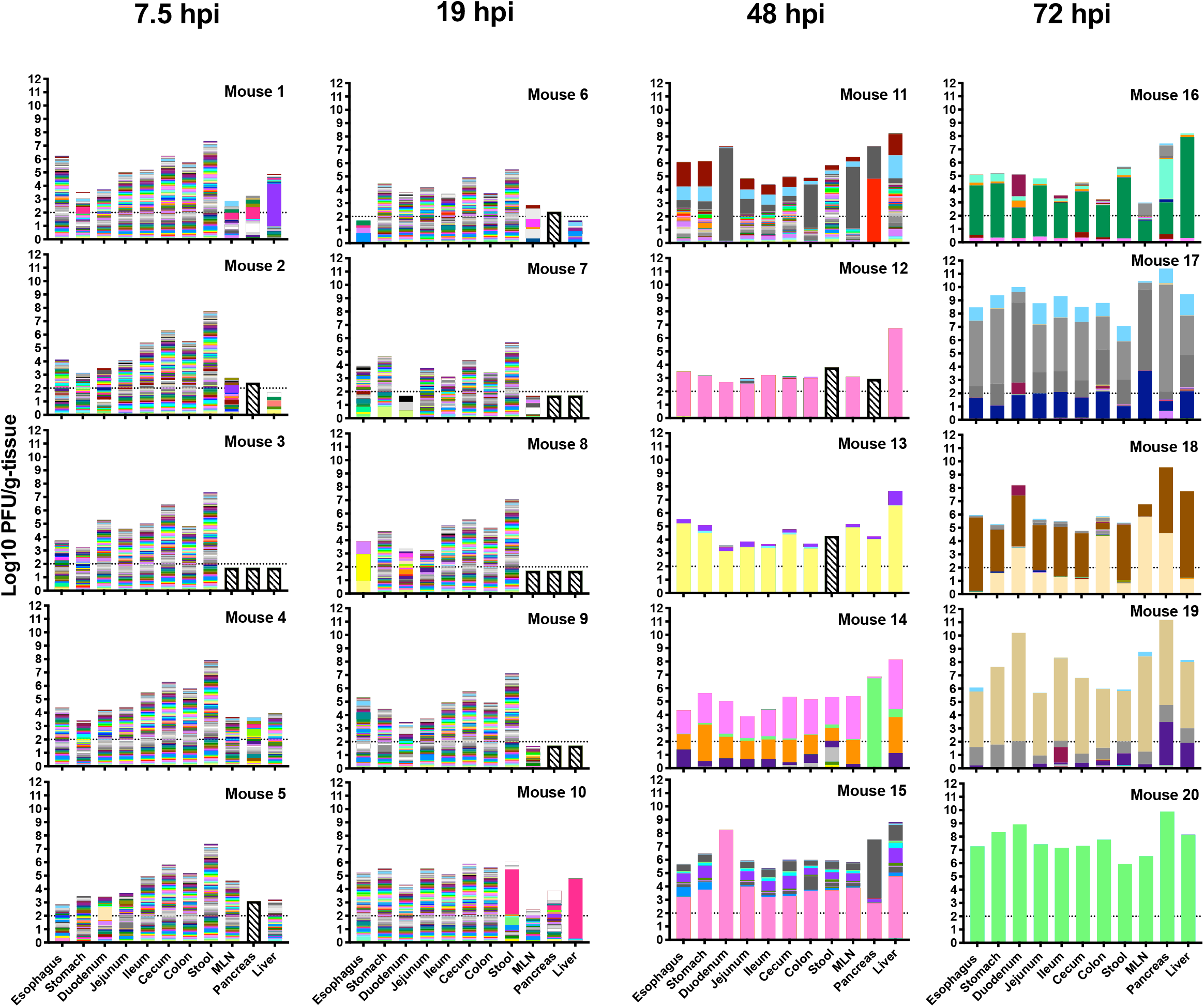
Virus population dynamics in gastrointestinal tract and extra-intestinal tissues over time. *Ifnar-/-* mice were orally infected with 1×10^9^ PFU of the CVB3 barcode library and were dissected 7.5 hpi, 19 hpi, 48 hpi, or 72 hpi. Height of bar corresponds to titer of virus, and the height of color bands is proportional to frequency of a viral barcode within sample, with total frequencies equaling 1. Horizontal dashed line is limit of detection for plaque assay, and bars with checkerboard pattern were below sequencing threshold.

To quantify viral population changes, we used two metrics of intra-sample diversity: 1) the total number of viral barcodes detected and 2) the Shannon diversity index, which is sensitive to the number of barcodes present as well the relative proportions of barcode frequencies. We analyzed viral diversity in GI tract tissues (esophagus, stomach, duodenum, jejunum, ileum, cecum, proximal colon, and stool) vs. extra-intestinal tissues (mesenteric lymph nodes (MLN), pancreas, and liver) to determine whether populations differed in tissues directly exposed to virus vs. tissues exposed following systemic spread. The total number of barcodes and Shannon diversity across the GI tract were high at 7.5 and 19 hpi (Supplementary Fig. 1). However, by 48 hpi and culminating at 72 hpi, the number of barcodes and Shannon diversity diminished across the GI tract. In contrast, the number of barcodes present and Shannon diversity were generally lower for extra-intestinal tissues regardless of time point. These results suggest broad early dissemination of the viral population in the GI tract and slightly reduced dissemination to extra-intestinal sites, with clearance and monopolization by a few pool members at all sites later in infection.

The broad “take over” by a few viruses in nearly all tissues observed at later time points could be due to adaptive mutations that confer a fitness advantage. To test if the high-frequency viruses harbored viral mutants that were selected for, we obtained CVB3 genome-wide consensus sequence for two tissues, MLN and liver. There were no detectable mutations in 5 of the 8 samples (Fig. 3). The remaining samples had 2 non-coding mutations (T7343C and C3217T) and 1 coding mutation (C6506T; 3D^H199Y^) (Fig. 3). Although consensus sequencing cannot detect low frequency mutations within a population, these results suggest that a major adaptive mutation does not drive the population monopolization effects we observe here.

**Figure 3.**
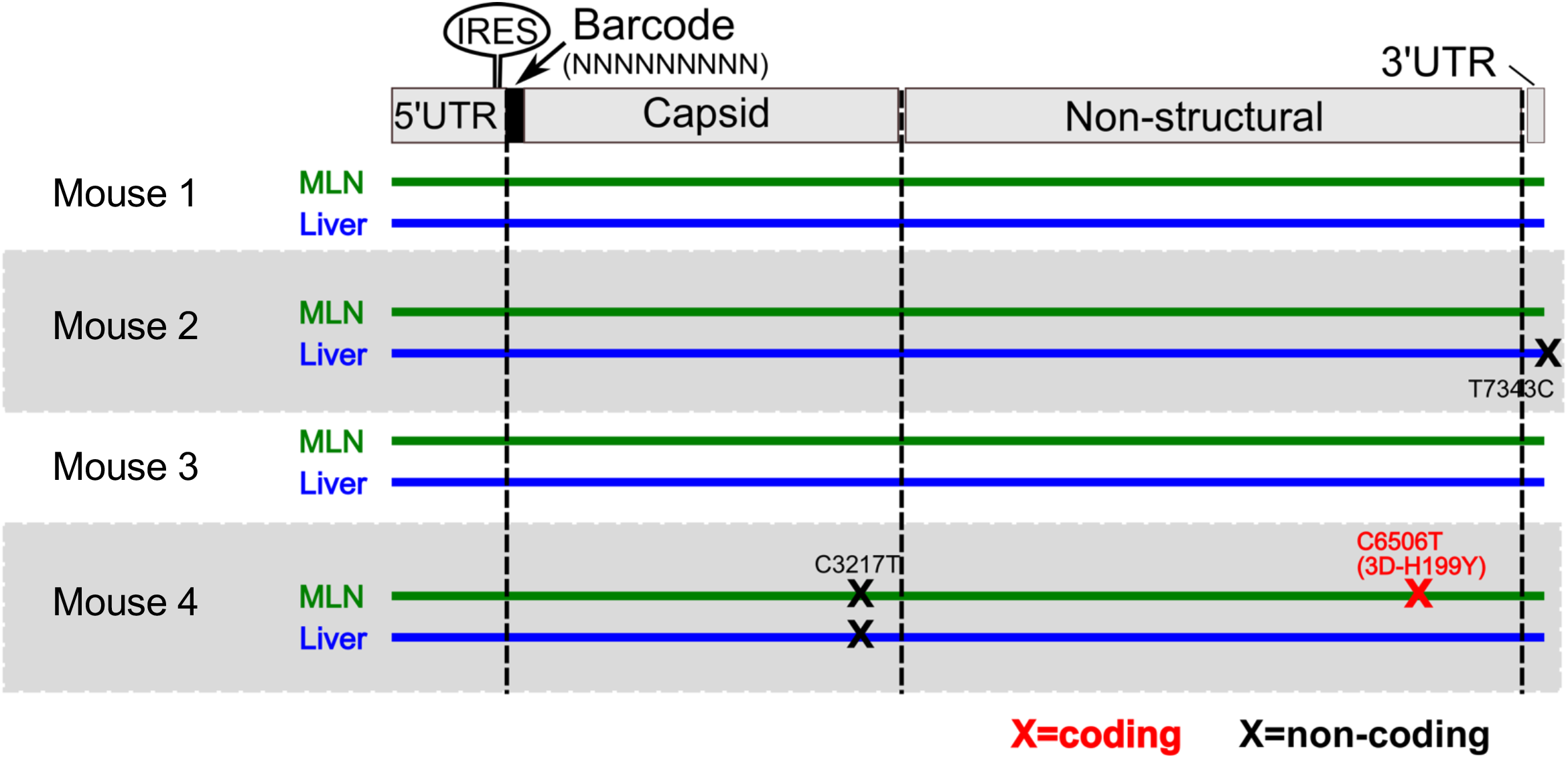
Monopolization of viral populations occurs in absence of detectable adaption. Sanger consensus sequencing of CVB3 whole genome was performed for 48 hpi MLN and liver samples from four representative mice. Each line represents the genome of consensus sequence derived from each sample, and an X represents a consensus mutation at that site.

### Multiple viruses replicate in all tissues prior to monopolization

One hypothesis for monopolization by viral subpopulations is that the small number of viruses detectable at later time points had undergone replication whereas the other population members detectable at early time points were unreplicated viruses from the original inoculum. To test this hypothesis, we used light-sensitive viruses to discriminate between unreplicated inoculum virus and replicated progeny virus. We have previously used light-sensitive, neutral red labeled barcoded virus to identify which barcoded viruses replicated in mice (18). Viruses propagated in the presence of neutral red incorporate the dye into capsids and lose infectivity when exposed to white light. However, upon replication in the dark, newly replicated viruses are light insensitive, thus facilitating discrimination inoculum from replicated virus. Using this strategy, we produced a neutral-red labeled stock of barcoded CVB3 viruses. Confirming light sensitivity, viral titer from this stock was reduced 250,000-fold after exposure to light. In the dark, we orally infected mice with 1×10^9^ PFU, harvested tissues, and processed tissues. Each tissue homogenate was split and half remained in the dark and the other half were exposed to light. We then performed experiments to 1) examine viral titers and the percentage of viruses that were leftover inoculum vs. replicated virus, and 2) quantify viral barcode population diversity among the inoculum vs. replicated virus subsets within each tissue (Fig. 4A).

**Figure 4.**
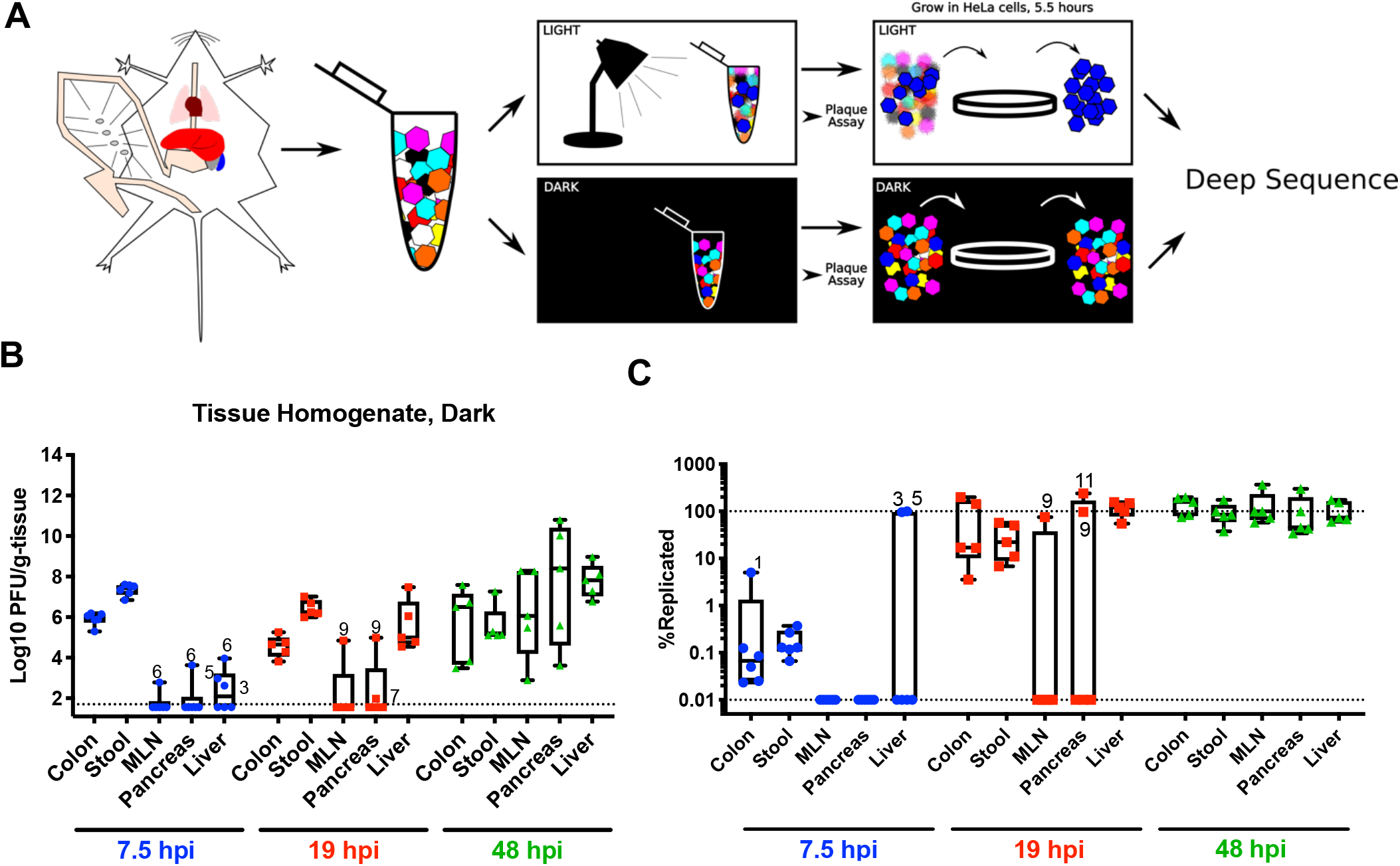
CVB3 replicates within extra-intestinal tissues by 19 hpi. A. Mice were orally infected with neutral red labeled CVB3 barcoded viruses in the dark. Mice were dissected in the dark and tissue samples were homogenized and either kept in the dark or exposed to light. Virus was quantified by plaque assay, then amplified for a single cycle in HeLa cells, and deep sequencing was performed on barcodes of progeny. In the example here, the blue virus had replicated *in vivo*, and therefore it was light insensitive and amplified in HeLa cells. B. Viral titers in samples kept in the dark, representing both input/inoculum and replicated viruses. Whiskers show max to min; points labeled with numbers indicate mouse number. C. Percent of replicated virus (PFU from light-exposed samples divided by PFU of dark-exposed samples multiplied by 100). Lower dotted line indicates limit of detection derived from plaque assay for light-exposed samples and upper dotted line indicates 100% replication within a population.

First, to determine total tissue titers and the percent of virus that had undergone replication, samples were quantified by plaque assay. Titers from samples kept in the dark, representing total tissue titers, are shown in Figure 4B. Trends were similar to data shown in Figure 2, with low early extra-intestinal tissue titers that increase with time. To reveal the percentage of viruses that had undergone replication, we divided the light titers by the dark titers for each tissue and multiplied by 100. As shown in Figure 4C, a fraction of virus in stool and colon was light insensitive at 7.5 hpi, indicating some viral replication had occurred *in vivo* even at this early time point (Fig. 4C). The amount of light-insensitive virus was 10-fold higher than the background of light insensitive/unlabeled virions in the inoculum, arguing that the majority of light-insensitive virus in these samples had replicated *in vivo* (Supplementary Fig. 2). The majority of virus in extra-intestinal tissues was still light-sensitive, indicating little viral replication had occurred by 7.5 hpi (Fig. 4C). At 19hpi, two mice had substantial viral replication in the MLN and pancreas, but nearly all virus in the liver had replicated *in vivo* (Fig. 4C). By 48 hpi, nearly all virus in all tissues was light insensitive (Fig. 4C).

Second, we quantified viral barcode diversity in each tissue, differentiating diversity in inoculum vs. replicated populations. Because RNA from neutral red inactivated virus could still be amplified by RT-PCR (data not shown), we enriched for viruses that had replicated *in vivo* by exposing to light and amplifying light insensitive viruses in HeLa cells for a single viral replication cycle (Fig. 4A). Viruses that had replicated *in vivo* would be insensitive to light exposure and undergo viral RNA amplification in HeLa cells, whereas inoculum viruses would not undergo RNA amplification. In parallel, the same procedure was used on samples kept in the dark, to amplify both inoculum and replicated viral RNAs. We then deep sequenced the viral barcodes to assess the diversity of viral barcodes from inoculum (dark samples) or replicated viruses (light samples) in each tissue. At 7.5 hpi, diversity of replicated viral barcodes was high in stool in all animals (Fig. 5, Supplementary Fig. 3). In contrast, few replicated viral barcodes were present in the colon. Given the striking difference of viral barcode diversity between replicated viruses in the colon and stool, it is likely virus replicated in numerous sites along the GI tract. In extra-intestinal tissues at 7.5 hpi, we sporadically detected replicated viral barcodes, and these had low diversity (Fig. 5, Supplementary Fig. 3). At 19 hpi in stool, all 135 barcodes viruses had replicated (Fig. 5, Supplementary Fig. 3) in most mice. Colon samples contained more diversity at 19 hpi than at 7.5 hpi, although the diversity of replicated virus was still much lower than the total number of barcodes present. At 48 hpi, we could detect replicated viral barcodes in all samples, and diversity in nearly all tissues was low. Consistent with Figure 2 data, populations across each mouse were very similar, supporting a model of whole mouse monopolization by a subpopulation of virus. Taken together, diversity among replicated viruses in the intestinal tract was high at 7.5 and 19 hpi, but low at 48 hpi, and diversity among replicated viruses was low at all time points in extra-intestinal tissues. Therefore, we conclude that loss of viral diversity at 48 hpi was not due to only a small fraction of virus replicating *in vivo*.

**Figure 5.**
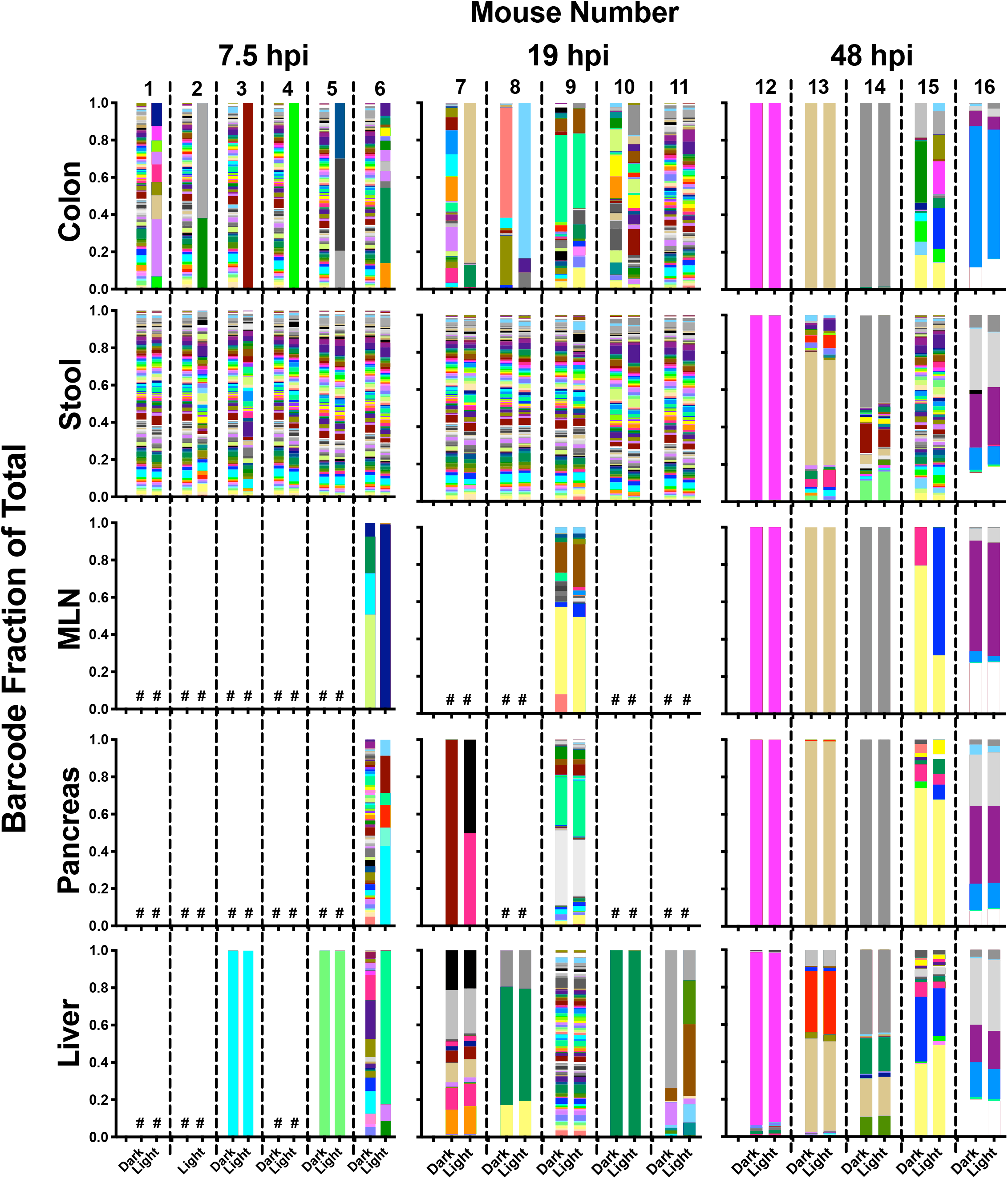
Diverse viruses within inoculum disseminate to extra-intestinal tissues and undergo replication. Ratio of viral barcodes within each sample, with viral barcodes derived from virus replicated *in vivo* (light-exposed) and total replicated + unreplicated inoculum viral barcodes (dark samples). The # symbol represents samples below sequencing depth threshold.

### Orally inoculated CVB3 rapidly spreads to extra-intestinal tissues in mice

Our data indicate that orally inoculated virus disseminates broadly within a few hours, but how quickly can virus disseminate to GI tract tissues and extra-intestinal sites? To assess how quickly and where oral virus inoculum disseminates before viral replication occurs, we infected *Ifnar-/-* and *Ifnar+/+* mice with 1×10^9^ PFU CVB3 and assessed viral titers at 20 minutes post infection. Surprisingly, we detected virus in the pancreas, liver, and MLN at this extremely early time point (Fig. 6A). Additionally, the early dissemination of CVB3 to extra-intestinal sites was observed in *Ifnar-/-* mice, but not in *Ifnar+/+* mice. In *Ifnar-/-* mice, viral barcode diversity was high in extra-intestinal tissues, indicating rapid dissemination of a relatively diverse population (Fig. 6B). To determine whether the rapid systemic dissemination was specific to CVB3, we quantified dissemination of related enteric virus, poliovirus. Poliovirus did not disseminate to extra-intestinal tissues by 20 minutes post infection (Fig. 6C), indicating rapid viral spread is not shared among all enteroviruses. Taken together, these data indicate that CVB3 can disseminate systemically very soon after oral infection.

**Figure 6.**
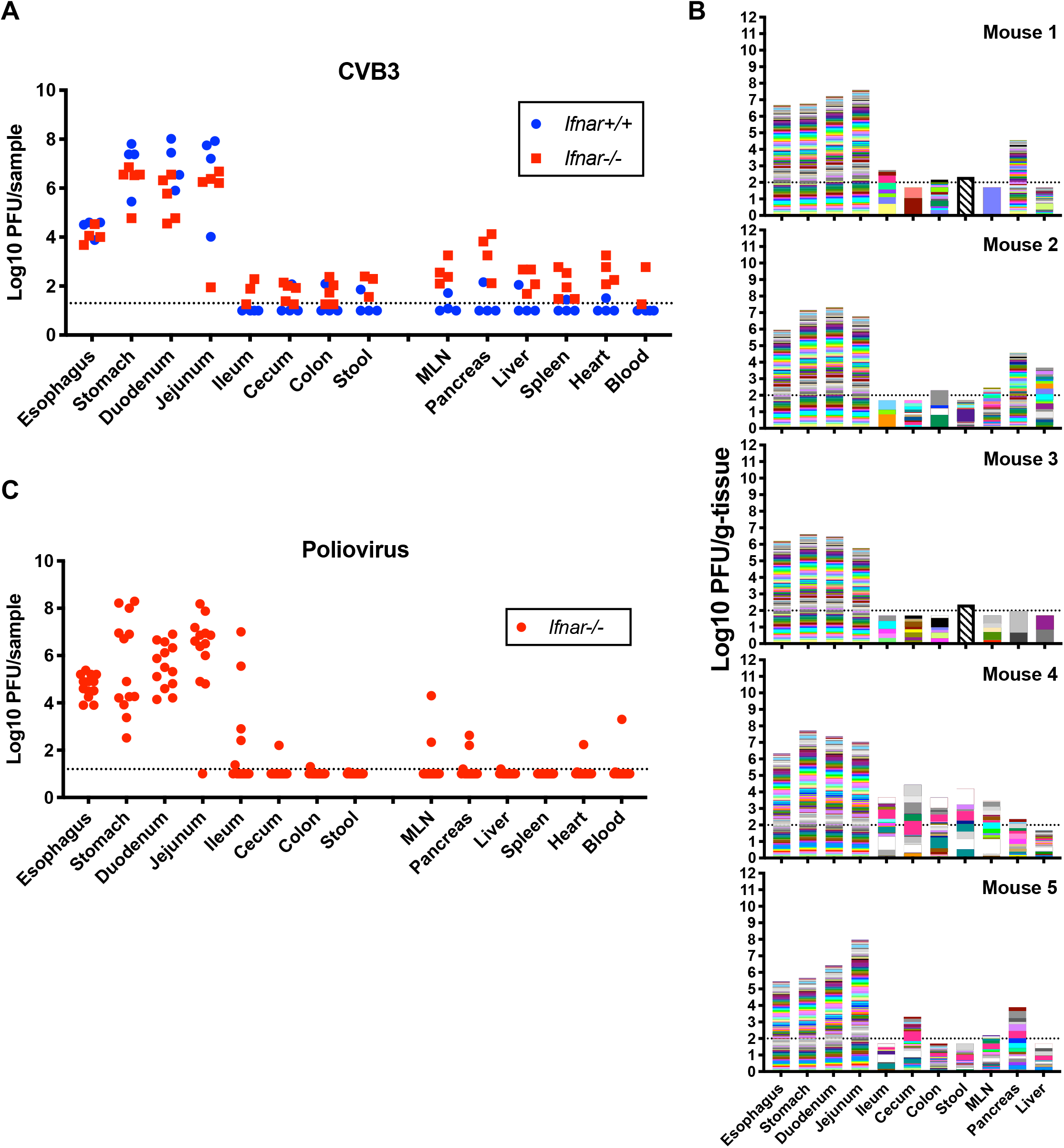
Orally inoculated CVB3 rapidly spreads to extra-intestinal tissues in mice. A. *Ifnar+/+* and *Ifnar-/-* mice were orally inoculated with 1 × 10^9^ PFU CVB3 and tissues were collected at 20 mpi. Virus was quantified in homogenized tissues by plaque assay (n=4). B. Dissemination of CVB3-barcode collection at 20 mpi (n=5). C. *Ifnar-/-* mice were orally inoculated with 1 × 10^9^ PFU poliovirus infected with poliovirus and tissues were collected at 20 mpi. Virus was quantified in homogenized tissues by plaque assay (n=13).

## Discussion

In this work, we engineered a barcoded collection of CVB3 to monitor viral populations across time and space in orally infected mice. Consistent with the our previous observation that the GI tract limits poliovirus diversity (6), we detected lower viral diversity in extra-intestinal tissues compared with the GI tract (Fig. 2). We also observed monopolization of viral populations by a small viral sub-population, and this viral sub-population was shared among virtually all tissues. In our earlier work with poliovirus, tissues were harvested upon disease onset—generally 72 hpi or later. At these late time points, we observed low poliovirus population diversity and we concluded that this loss of diversity was due to combined bottlenecks from mouth-to-gut and gut-to-blood (6). However, here we found that replication of diverse CVB3 populations in both the GI tract and extra-intestinal tissues preceded loss of diversity (Fig. 5). Thus, loss of CVB3 diversity was not only due to a small subset of viruses from the inoculum replicating *in vivo*. Strikingly, nearly all tissues in each mouse were monopolized by a small sub-population of viral barcodes (Fig. 2, 5). These results suggest that viral populations in separate tissues disseminated and intermingled, and that the viral population in a given site could be subsumed by an invading viral population.

Due to the propensity for RNA viruses to rapidly develop diverse populations, it was possible the monopolized populations we observed were the consequence of mutations that were selected for and enriched. Indeed, others have reported the selection for poliovirus mutations uniquely adapted for specific tissues (19). However, by 48 hpi, we found that nearly all viral populations within a mouse were overwhelmed by the same small viral sub-population. In this time frame, we did not find detectable mutations within the whole genome consensus sequences that could explain adaptive advantage of this sub-population (Fig. 3). In the absence of major adaptive mutations, a key future question is to define 1) how viral subpopulations dominate within a tissue and whole animal, 2) the anatomic source of monopolizing viral population, and 3) whether clearance enables monopolization.

In order to delineate the population dynamics of an enteric virus over time, we set out to measure kinetics of CVB3 dissemination after oral feeding of virus. We found that virus could reach extra-intestinal tissues as soon as 20 minutes post-infection in *Ifnar-/-* mice, but not in *Ifnar+/+* mice (Fig. 6). Previous work has shown that interferon gamma can influence GI barrier permeability, but roles for interferon alpha/beta in rapid viral dissemination have not been described (20). Furthermore, poliovirus did not disseminate to extra-intestinal sites by 20 minutes post-infection, indicating a unique rapid dissemination mechanism for CVB3 that is not shared for all enteroviruses. What mechanisms could explain this rapid dissemination? Either passage across the intestinal epithelia or via M cells can lead to viral access to the lymphatic system, then to the portal vein, and subsequently the bloodstream, or other mechanisms may contribute. Future experiments will explore the physiological routes of rapid enterovirus dissemination and the contribution of type 1 interferon signaling to GI permeability.

In conclusion, we find that diverse viral populations disseminate and replicate in GI and extra-intestinal sites, but viral population diversity is severely restricted post-replication. These results support a model of early broad dissemination and subsequent clearance for an enteric virus.

## Materials and Methods

### Cells and Media

293T and HeLa cells were maintained in Dulbecco’s modified Eagle’s medium (DMEM) with either 10% normal calf serum (HeLa) or fetal calf serum (293T), 2 mM L-glutamine, and 10 mM HEPES. Cells were maintained at 37°C and 5% CO_2_. All transfections with viral plasmids were performed with Lipofectamine 2000 (ThermoFisher). Our HeLa cells (originally from Karla Kirkegaard’s lab) are highly permissive for replication of enteroviruses compared with other HeLa cell lines including the ATCC line.

### Viruses

In this work, we used CVB3-H3 strain (Woodruff U57056.1, with 16 nt differences of unknown origin), and poliovirus-Mahoney serotype 1. All poliovirus work was done in WHO-approved elevated BSL2+/poliovirus conditions. Stocks were generated by transfecting 293T cells with 1) plasmids containing viral genomes under T7 promoter, and 2) plasmid expressing T7 polymerase. For the CVB3 barcode viruses, each individual stock was propagated separately to control the ratio of each virus in the final mixture. After 2.5 days, cells were freeze-thawed 3 times, and clarified supernatant was added to HeLa cells. Cells were incubated until cytopathic effects were evident, and were then freeze-thawed 3 times. These initial viral stocks were quantified by plaque assay and then amplified in HeLa cells at multiplicity of infection >10. When cytopathic effects were observed, cells were collected by scraping, freeze thawed 3 times, and resuspended in a small volume of PBS containing 0.1 mg/ml MgCl_2_ and CaCl_2_. Virus within clarified supernatant was quantified and used for experiments.

### Plaque Assay

Samples were serially diluted, and 200 ul was plated onto HeLa cells in 6 well plates (70% confluent). Cells were incubated at room temperature >20 minutes, and media with 1% agar were added to wells, and incubated at 37°C for 2-3 days. Wells were stained with 1% crystal violet in 20% ethanol, rinsed with water, and plaques counted.

### Neutral Red/Light Sensitive Viruses

Virus stocks were amplified as above, except in the presence of 50 μg/mL neutral red (6). Neutral red labeled viruses were handled in the presence of a red light to maintain infectivity. To quantify percent light-sensitive (inoculum/unreplicated) virus, an aliquot of virus was exposed to white light from a desk lamp for 20 min. For tissue homogenates, samples were exposed to light for 1 h with occasional mixing.

### Virus Growth Curve

HeLa cells were infected in suspension at MOI 10 for 30 min on ice, washed 3 times, and 50,000 cells were plated per well in 96 well plate. At indicated time points, plates were frozen. Three replicates were performed for each condition at each time point, and the experiment was repeated 3 times. For the growth curve, virus stock from a single representative barcode was used.

### Construction of Virus Barcode Library

To insert the 9 nt barcode, primers (IDT) were designed (NEBaseChanger) with random overhangs on the 5’ end to be inserted between nucleotides 708 and 709 of the CVB3 genome. PCR was performed using Q5 high-fidelity DNA polymerase (NEB, M0491L). PCR product was treated using the KLD Enzyme Mix (NEB, M0554S), and transformed into competent bacteria (Zymo, T3007). Individual colonies were screened for the presence of the 9 nt barcode by colony PCR and then verified by Sanger sequencing. Individual stocks of virus were prepared as described above, mixed at equal PFU, then concentrated by ultracentrifugation (66,549 x g overnight through a 10 ml 30% sucrose).

### Mice

All mice experiments were approved by the UT Southwestern Institutional Animal Care and Use Committee. C57BL/6J, C57BL/6-*Ifnar-/-*, C57BL/6-PVR (transgenic for the human poliovirus receptor), and C57BL/6-PVR-*Ifnar-/-* (transgenic for the human poliovirus receptor) (15) mouse strains were maintained using standard animal husbandry practice in UTSW animal facilities.

### Dissections and Tissue Processing

Tissues were collected and snap-frozen on liquid nitrogen. After weighing, beads (Fisher Scientific, NC9862662) and PBS with 0.1 mg/ml MgCl_2_ and CaCl_2_ were added, and were homogenized (Next Advance Bullet Blender Storm). Samples were divided into portions for RNA isolation and viral quantification. Fractions destined for plaque assay were freeze thawed 3 times, spun at 16,000 x g, and virus was quantified within the supernatant. RNA was isolated from intestinal contents and stool using Viral RNA Extraction Kit (Zymo, R1035), and from tissues using TRI LS (Sigma Aldrich, T3934), per manufacture’s protocol.

### Sequencing and Analysis of Barcode Library

cDNA was synthesized using primers and Superscript II enzyme (Fisher Scientific, 18064071). Libraries were produced by amplifying cDNA using primers targeting the barcoded viral locus in PCR. Primers contained Illumina adaptor sequences, Illumina indices within reverse primers for demultiplexing by the sequencing core, and custom indices within forward primers to increase number of samples to multiplex and to phase the sequencing reaction. Samples were gel purified and pooled by amplicon band intensity. These pools were combined together, normalizing by nucleic acid concentration. Sequencing was performed on Nextseq Illumina sequencer with 75 cycles at UTSW McDermott Sequencing Core. FASTQ files were demultiplexed by reverse index, then demultiplexed by the forward index. Barcodes were extracted from reads plus surrounding 30 nucleotides of viral genomic sequence to ensure barcodes were associated with virus sequence. Barcode counts were enumerated with the “grep” command line function, with each barcode sequence serving as a search string. Pilot studies indicated sequencing depth of at least 1000 reads/sample was sufficient to recapitulate barcode ratios; we set this as the threshold. Because of variable sequencing depth, we set 0.0005 viral barcode reads/total reads-sample as the cutoff to count a barcode as present.

## Graphing and Statistics

All graphs and statistics were produced with Graphpad Prism. Shannon diversity index was calculated using the R package “vegan” (21). Figures were prepared using Inkscape.

## Acknowledgements

We thank members of the Pfeiffer lab for helpful discussions. This work was supported by a Burroughs Wellcome Fund Investigator in the Pathogenesis of Infectious Diseases award (JKP) and R01 AI74668 (JKP). JKP is a Howard Hughes Medical Institute Faculty Scholar. MRL was supported in part by NIH grant T32 GM109776. BTM was supported by NIH grant T32 AI007520 and F32 AI138392.

---

**Supplemental Figure 1 (Related to Figure 2). Viral barcode diversity decreases over time and is lower in extra-intestinal tissues.**

Comparing changes in diversity over time, we quantified the total number of barcodes detected (A) and Shannon diversity (B) for GI tract or extra-intestinal tissues (MLN, pancreas, and liver) combined. Data in B were analyzed by Kruskal-Wallis test with Dunn’s correction for multiple comparison; only significant results are shown, * *p*<0.05. To directly compare GI vs. extra-intestinal sites, we analyzed the number of barcodes detected (C) and Shannon diversity (D). For data in D, we performed mixed effects analysis with Sidak’s correction for multiple comparison; only significant results are shown.

**Supplemental Figure 2 (Related to Figure 4). Nearly all light-insensitive virus recovered from mice is higher than background in neutral red labeled virus stock.** A small fraction of viruses within a neutral red stock are light insensitive, which can confound interpretation of results from subsequent mouse experiments. To rule this out, we quantified the fraction of light insensitive viruses in our stock and compared it with the data from mouse experiments. We found that 1/4000 PFU in neutral-red labeled virus inoculum was light-insensitive. Assuming 1) neutral-red labeled and unlabeled virions equally penetrated each tissue, and 2) no viral replication occurred, then 1/4000 PFUs from dark-treated tissues would be light insensitive inoculum PFUs. This sets the background level for each sample. The light-treated virus titer was divided by this background level to obtain the fold signal over background. Anything over 1 indicates virus replication occurred, but probability for *bona fide* viral replication increases exponentially up the Y-axis. Numbers indicate mouse number.

**Supplemental Figure 3 (Related to Figure 5). Viral barcode diversity from viruses replicated *in vivo*.**

Number of viral barcodes in each tissue at each time point from samples exposed to light and amplified for a single cycle in HeLa cells.

A. Number of barcodes present of replicated virus in each sample.

B. Shannon diversity (H’) of replicated viral barcodes. 2-way ANOVA, Tukey multiple comparison; only significant results are shown, * *p*<0.05.

